# Definition of the effector landscape across 13 Phytoplasma proteomes with LEAPH and EffectorComb

**DOI:** 10.1101/2023.12.06.570357

**Authors:** Giulia Calia, Alessandro Cestaro, Hannes Schuler, Katrin Janik, Claudio Donati, Mirko Moser, Silvia Bottini

## Abstract

**Background:** Crop pathogens are a major threat to plants’ health, reducing the yield and quality of agricultural production. Among them, the *Candidatus* Phytoplasma genus, a group of fastidious phloem-restricted bacteria, can parasite a wide variety of both ornamental and agro-economically important plants. Several aspects of the interaction with the plant host are still unclear but it was discovered that phytoplasmas secrete certain proteins (effectors) responsible for the symptoms associated with the disease. Identifying and characterizing these proteins is of prime importance for globally improving plant health in an environmentally friendly context.

**Results:** We challenged the identification of phytoplasma’s effectors by developing LEAPH, a novel machine-learning ensemble predictor for phytoplasmas pathogenicity proteins. The prediction core is composed of four models: Random Forest, XGBoost, Gaussian, and Multinomial Naive Bayes. The consensus prediction is achieved by a novel consensus prediction score. LEAPH was trained on 479 proteins from 53 phytoplasmas species, described by 30 features accounting for the biological complexity of these protein sequences. LEAPH achieved 97.49% accuracy, 95.26% precision, and 98.37% recall, ensuring a low false-positive rate and outperforming available state-of-the-art methods for putative effector prediction. The application of LEAPH to 13 phytoplasma proteomes yields a comprehensive landscape of 2089 putative pathogenicity proteins. We identified three classes of these proteins according to different secretion models: “classical”, presenting a signal peptide, “classically-like” and “non-classical”, lacking the canonical secretion signal. Importantly, LEAPH was able to identify 15 out of 17 known experimentally validated effectors belonging to the three classes. Furthermore, to help the selection of novel candidates for biological validation, we applied the Self-Organizing Maps algorithm and developed a shiny app called EffectorComb. Both tools would be a valuable resource to improve our understanding of effectors in plant–phytoplasmas interactions.

**Conclusions:** LEAPH and EffectorComb app can be used to boost the characterization of putative effectors at both computational and experimental levels and can be employed in other phytopathological models. Both tools are available at https://github.com/Plant-Net/LEAPH-EffectorComb.git.

## Background

Plants live in a constantly changing environment and have developed high phenotypic plasticity including rapid responses and adaptations to environmental factors (1,2). In the case of parasitism, interactions are based on a molecular dialogue between the pathogen and its host. Plants and parasites are engaged in an arms race where the plant develops stratagems to detect the bio-aggressor while the latter aims to bypass the host’s immune system (3–6). Parasites modulate the immune response, cell signaling, metabolism or even the development of the plant by acting on its transcriptome regulation (7–9). This modulation of gene expression relies on the secretion of specific pathogenicity factors called effector proteins. Effectors’ arsenal and their delivery system vary from species to species although they have the common aim to interfere with the host metabolism to their advantage, and to hamper the immune system, enhancing the pathogen survival and the development of the disease (10–13). Advancing our knowledge about the functioning and success of biotic interactions is of prime importance towards improving plant health globally in an environmentally friendly context.

The increasing availability of genome sequences of plant pathogens allowed dramatic advances in the characterization of pathogenicity mechanisms and the development of tools to improve effector prediction. For instance, EffectorO is a machine learning model exclusively trained on the N-terminal of oomycetes effector proteins (14). Another example is EffectorP1.0 to 3.0, an ensemble of machine learning models to predict both effectors and their localization (apoplastic or cytoplasmic) trained on both fungi and oomycetes to search for characteristic enrichment of amino acids and their properties (15–17). Deepredeff is a convolutional neural network trained on bacteria (gram+ and gram-), fungi and oomycetes sequences (18). DeepT3 combines different deep-learning algorithms to predict effectors secreted by the bacterial type III secretion systems (T3SSs) in Gram-(19). Finally, EffectiveT3 is a machine learning model to predict whether effector proteins are secreted by T3SS (20). Despite these tools allowed to advance effector knowledge for some plant pathogens, there is still a lack of prediction methods for several pests hampering a global characterization of pathogenic mechanisms.

A clear example of this category is the “*Candidatus* Phytoplasma’’ genus, a group of plant-pathogenic, phloem-restricted, bacteria assigned to the Mollicutes class, cell wall-less and pleomorphic bacteria, ranging from 0,2 to 0,8 μm in size (21–24). Phytoplasmas are causal agents of diseases in a large number of crops, ornamental plants and trees that result in plants altered development and huge yield losses (25–28). Characteristic symptoms of the disease are witches’ broom agglomerates of young branches, increased proliferation of shoots, yellowing of the leaves, virescence (flower organs become greeny), phyllody (flowers develop into leaf-like flowers), dwarfism, reduced in size and tasteless fruits (23,29,30). The transmission from plant to plant occurs by insects, which become vectors of the pathogen once they acquire the phytoplasma during phloem-feeding (31–33). The parasite behaviour of phytoplasmas and the consequent impossibility of cultivating them under *in vitro* conditions have hindered experimental studies focusing on the identification of effector proteins. Similarly, *in silico* identification of effector proteins exclusively based on the overall sequence similarities is inefficient due to their amino acid sequence variability. As a consequence, to date, very few effector proteins have been identified in phytoplasmas (34). TENGU, a small secreted peptide encoded by onion yellow phytoplasma, causes dwarfism and altered flower structures (35). SAP05 which interferes with plant vegetative growth is a secreted protein found from ‘*Ca*. Phytoplasma asteris’ (36,37). SAP11-like proteins, first identified from aster yellow witches broom phytoplasma, cause abnormal proliferation of young shoots and changes in leaf shape (36,38). SAP54 and PHYL1 are two homologous effectors that belong to the phyllogen family and cause flower malformations including phyllody, virescence and proliferation (36,39,40). The identification of further effector proteins would contribute to a better understanding of the biological mechanisms involved in disease development, providing new knowledge for the development of novel intervention strategies for pest management.

Nowadays, current methods for effector identification in phytoplasmas are only based on the presence of signal peptide, fairly reconstructing phytoplasmas’ secretome. However, it is important to consider that not all the secreted proteins are effectors and that some effector proteins are secreted by nonclassical pathways (41,42). Moreover, it is shown that in Gram+ bacteria, ancestors of phytoplasmas, the hydrophobic region of signal peptides is longer than usual, making them more similar to transmembrane regions and undetectable by software specifically designed for signal peptide prediction (43). Thus, recent methods rely on the combined predictions by software designed for both signal peptide and transmembrane domain detection (43,44), leading a large number of candidates to be tested without any prioritization assessment. Therefore, there is an urgent need for a tailored method to predict and characterize phytoplasmas effectors.

To address these needs, we developed LEAPH (ensemb**L**e model for **E**ffector cl**A**ssification in **PH**ytoplasmas) a computational method composed of an ensemble of four supervised learning models to capture distinct sets of features and efficiently predict putative disease-related proteins in phytoplasmas. Indeed, LEAPH predicts effectors with ∼97% accuracy outperforming existing prediction methodologies tailored for other pathogens. Our ensemble model was trained on 479 proteins coming from more than 50 phytoplasma species and then applied, as a use-case scenario, on 13 proteomes (ranging from 327 to 730 proteins), allowing us to identify a comprehensive landscape of putative candidate effectors. We used the Self-Organizing Map algorithm to describe the properties of this landscape, which combines clustering and dimensionality reduction to embed the protein sequences similarities based on the feature profiles. Therefore, neighboring points on the map represent proteins sharing similar features providing the first reference map for phytoplasma effectors. We developed a user-friendly shiny application, called EffectorComb, to investigate the effector protein map. Overall, LEAPH and EffectorComb offer the possibility to predict, interpret, and explore the resulting protein candidates, thus boosting the experimental validation process from the very beginning.

## Materials and Methods

### Training Datasets

For the training dataset sequences selection and curation, we used the UniProt database (release 2022_01) (45), selecting proteins uniquely belonging to the phytoplasmas TAXID 33926.

### Positive dataset

The positive dataset is created using two filters. Firstly the field “Protein Names” is searched using the AND operator and each of the following regular expressions: “*Effector*”, “*TENGU*”, “*SAP54*”, “*SAP11*”, “*SAP05*”, “*PHYL1*”. Secondly, the resulting list is pruned from records that have the terms “putative” or “fragment” in the “Protein names” field. The result is a set of 174 proteins from 53 phytoplasmas species (Figure 1a). Other 10 experimentally validated proteins are retrieved from A. Hoshi et al., 2009 (35), TENGU, X. Bai et al., 2008 (36), SAP54, X. Bai et al., 2008, M. Kube et al., 2009 (36,46), SAP11, X. Bai et al., 2008 (36), SAP05, K. Maejima et al., 2014 (40), PHYL1. In total, the positive dataset accounts for 184 protein sequences (Additional Table 1.1).

**Figure 1:**
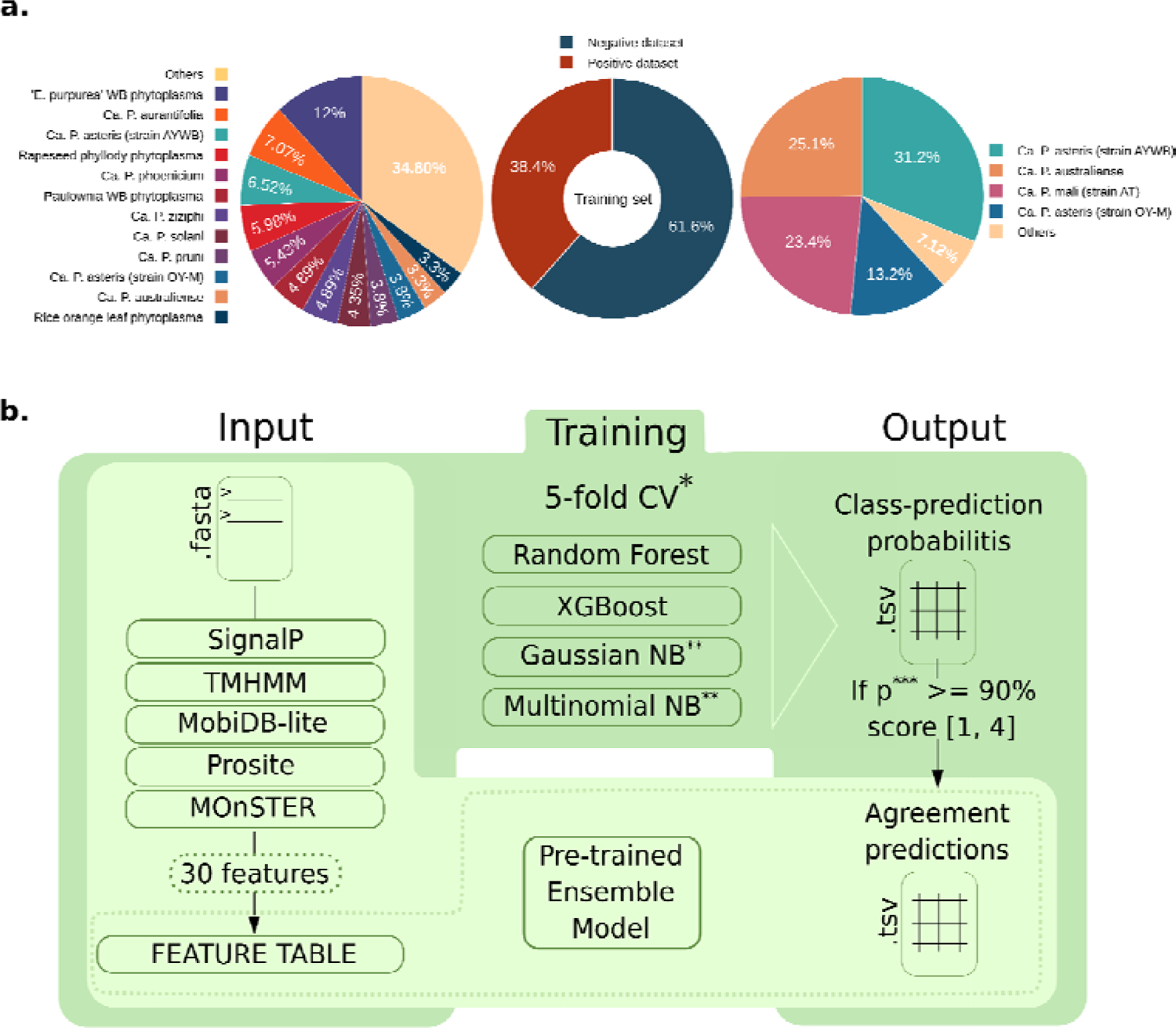
Training dataset composition and LEAPH workflow. (a) training dataset composition of positive and negative set proportion (in the center) and phytoplasma species for the positive dataset (on the left) and negative dataset (on the right). Refer to Additional Table 1.1 and 1.2 for details. **(b)** LEAPH workflow. The dark green box highlights the training process while the light green box represents the application process of LEAPH; the dashed box depicts the particular use case in which a feature table is already available, thus ready to be used as input for the pre-trained LEAPH model.

### Negative dataset

The negative dataset is constructed by manual inspection of UniProt database entries having evidence for revision in “Protein existence” field AND having function not related to the known effectors’ activity (e.g. no “*SVM*” or “*trigger factor*” in “Protein Name” field). The remaining list is checked for the absence of a cross-reference with the PHI-base database (47,48), meaning that there is no known interaction between these proteins and the host ones. The negative dataset is finally composed of 295 non-effector proteins from 12 phytoplasma species (Additional Table 1.2, Figure 1a).

### Proteomes Datasets

We used LEAPH to predict putative effector proteins in 13 phytoplasma proteomes: ‘*Ca.* P. asteris’ (strain AYWB-UP000001934 and OY-UP000002523), ‘*Ca.* P. australiense’ (UP000008323), ‘*Ca.* P. mali’ (strain AT-UP000002020), ‘*Ca.* P. oryzae’ (strain Mbita1-UP000070069 and S10-UP00024934), ‘*Ca.* P. phoenicium’ (strain ChiP-UP00023867 and SA213-UP00003708), ‘*Ca.* P. pruni’ (strain CX-UP00003738), ‘*Ca.* P. tritici’ (2568526557 from JGI-GOLD (49)), ‘*Ca.* P. vitis’ (CP097583.1 from NCBI-GeneBank (50)), ‘*Ca. P. ziziphi*’ (CP091835.1 from NCBI-GeneBank) and ‘Maize bushy stunt phytoplasma’ (CP015149.1 from NCBI-GeneBank]). The proteomes are retrieved from Uniprot If not indicated otherwise. The main characteristics of the 13 proteomes are summarized in Additional Table 2.

### LEAPH method description

LEAPH workflow consists of four main steps described hereafter and shown in Figure 1b. Briefly, in step 1, LEAPH takes as input the amino acid sequences of interest in FASTA format and computes the 30 features, described in the following paragraph, yielding as output the feature table. During the training process (step 2), the feature table is used as input for four classifier models to be trained. After a 5-fold cross-validation process, each of the best models is used to finally assign a class probability to every protein in the overall training dataset (step 3). LEAPH predicts a protein to be putative pathogenic if at least one of the four models gives a class probability higher than 90% and thereby assigns a score ranging from 1 to 4 to each protein based on the models’ agreement (step 4). The pre-trained model can be used to predict putative pathogenicity proteins in any set of protein sequences producing an output table that contains the resulting classification along with model agreement, mean prediction probability and the corresponding amino acid sequence for each protein.

### Features Calculation

The first step of LEAPH is the annotation of protein sequences and the extraction of features to describe protein sequence properties. We included a total of 30 features such as: (i) sequence length, (ii) signal peptide presence (using the D-score of SignalP 4.1 software (51) configured as in the work of Garcion et. al (43)) and transmembrane domain (TM) presence (TMHMM2.0 (52)). Specifically, we included different aspects of the transmembrane regions prediction, namely (iii) the number of predicted TMs, (iv) the expected number of amino acids in TMs, (v) the number of expected amino acids in the first 60 positions of TM helices, (vi) the probability that the N-terminal of the protein is in the cytoplasm, and (vii) a warning for possible misprediction of the TM-regions (when feature vi > 10), representing a potential signal peptide region (mTMR). Since effector proteins can be composed of intrinsically disordered regions (IDRs) (53) we also predicted (viii) the eventual presence and the length of IDRs (MobiDB-Lite3.0 (54)). Finally, we added 22 features concerning putative characteristic motifs of effectors: (ix-xx) by using Prosite1.86 (55), we calculated the occurrences of functional protein motifs belonging to the positive dataset; (xxi-xxx) by using MOnSTER (56) we obtained cluster of Prosite motifs (CLUMPs) considering their physicochemical characteristics. From MOnSTER application, we extracted six features representing the occurrence of each selected CLUMPs and four additional features, namely the occurrence of CLUMPs in 4 consecutive bins of 25% of the protein sequence. The configuration parameters for all used tools to obtain the 30 features and the main descriptions are indicated in Additional Table 3.

### Models

LEAPH is based on an ensemble learning approach that uses four classification algorithms: Random Forest (57) (RandomForestClassifier() from sklearn.ensemble), XGBoost (58) (XGBClassifier() from xgboost python library), Gaussian Naive Bayes (59) (GaussianNB() from sklearn.naive_bayes), and Multinomial Naive Bayes (60) (MultinomialNB() from sklearn.naive_bayes). The version of scikit-learn library is 1.1.1 for all the methods (61). See the following section and Additional Table 4 for more information on usage and parameter settings.

### Cross-validation and hyperparameter tuning

Because of the size and the unbalanced distribution of proteins between classes in the training set, we applied the stratified 5-fold cross-validation method (StratifiedKFold from scikit-learn 1.1.1). Each model is trained and tested five times on a different subset of the training data respecting class proportion (80% of data for the training set and the remaining 20% for non-overlapping test sets).

We performed a hyperparameter tuning via GridSearchCV from scikit-learn 1.1.1, for each fold, and for each classifier except for Gaussian Naive Bayes which was trained with the default parameters. With the grid search, the parameters are tested from a specified set of values. The starting parameters used in the training process, which differ from the default ones, are included in Additional Table 4. Altogether cross-validation and hyperparameter tuning allow to select the best model for each classifier algorithm to be used to predict new putative pathogenicity proteins.

To support the reproducibility of the machine learning method of this study, the machine learning summary table (Additional Table 4) is included in the supporting information as per DOME recommendations (62).

### Classification method

The best model for each classifier obtained from the previous phase is used to finally assign a class probability to every protein in the overall training dataset: if for at least one model the class probability is >= 90%, then the protein is considered as putative pathogenic. LEAPH assigns a score ranging from 1 to 4 to each protein based on the models’ agreement. The pre-trained model can be used to predict novel putative pathogenicity proteins in any set of proteins yielding an output table containing the classification prediction, the model agreement, the mean prediction probability and the corresponding amino acid sequence for each protein.

### Performance measures

To assess the performance of the prediction models used in this study, we used the following metrics: accuracy, precision, recall, and f1-score defined as:

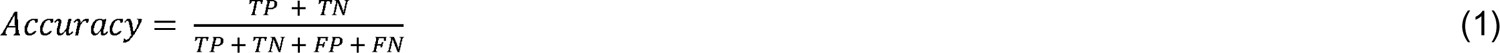

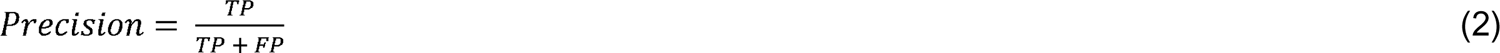

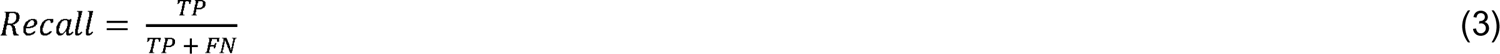

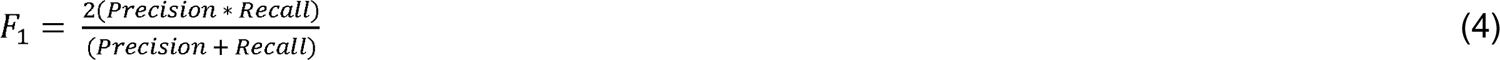

Where TP, TN, FP, and FN refer to True Positives, True Negatives, False Positives, and False Negatives, respectively. These measures are calculated using the sklearn.metrics library.

### Benchmark of effector prediction tools

The state-of-the-art method commonly used in phytoplasmas’ effector prediction consists of a combination of two predictors: SignalP4.1 and TMHMM2.0. Specifically, a protein is predicted as putative pathogenic if both signal peptide and mTMR are predicted. This method is included in our benchmark as the reference method for phytoplasmas effectors. We also compared the performance of LEAPH with other effector prediction tools on the training set. We considered EffectorP3.0 (17), which is mainly suitable for fungi and oomycetes putative pathogenicity protein, EffectorO (14) tested for oomycetes putative effector prediction and Deepredeff (18) used for both bacteria, fungi and oomycetes putative pathogenicity protein prediction. All the tools are used with default parameters or, where possible, in a configuration suitable for bacteria.

### Feature importance calculation

We assessed the feature importance for each classification algorithm using SHapely Additive exPlanations (SHAP) approach (63). SHAP is widely used for explainable machine learning and gives a more comparable spectrum of feature importance across different models. It measures the contribution of each feature to the final output using game theory concepts and feature permutations, assigning a SHAP value to each feature. Compared with the classical feature importance measurements, this additive feature method relies on maintaining the accuracy of the model, dealing with missing features and consistency in model-changing for the same data.

### SecretomeP2.0

To predict non-classically secreted proteins we used SecretomeP2.0 (64). Since the Gram-positive bacteria model is currently unavailable for the latest version of the tool, we used the Gram-negative bacteria one. We split the sequences into 100 sequences per fasta file (using seqkit split (65) and setting the parameter “-s 100”) to use the web server, and we cross-checked each prediction with the presence of a signal peptide predicted by both SignalP4.1 and TMHMM2.0, as SecretomeP2.0 web-service recommendations.

### Exploratory & Functional Analysis

We performed a Principal Component Analysis (PCA) on scaled values (MinMaxScaler from scikit-learn 1.1.1) of the 30 protein features described in the paragraph “***Features Calculation”***, and InterProScan5 (66) to predict protein domains associated with a known biological function on the predicted putative pathogenicity proteins identified by LEAPH. We then explored the differences in protein domains that occur in at least 1% of the sequences, according to the secretion modality of the predicted proteins.

### Effector proteins map

We used Self-Organizing Maps (SOM) (67) to exploit the properties of the putative pathogenicity proteins identified by LEAPH from 13 phytoplasmas proteomes. To build up the SOM we firstly scaled the values of each feature (MinMaxScaler from scikit-learn 1.1.1), excluding mTMR as a categorical, thus not suitable, feature and then used the aweSOM R package, which outputs a dynamic map. Through SOM visualization, points on the map in the same hexagonal cell share a similar feature vector and thus are indeed similar to each other. To find the optimal number of hexagonal cells to create the map, we tried different sizes of lattice grids from 8×8 up to 11×11, where we stopped because the 10×10 achieved the best balance between the errors (Quantization, Topographic and Kaski-Lagus) minimization and variance explained (see Additional Table 5 for further details).

### EffectorComb Shiny App

We developed an interactive Shiny App to investigate the obtained SOM. The app allows the user to retrieve a set of pathogenicity proteins predicted by LEAPH with properties of interest along with their sequence identifier and amino acid composition. It is also possible to download SOM interactive images .html files. EffectorComb can be also used to project new proteins, predicted by LEAPH application, on the pre-obtained SOM for a deep result exploration and comparison.

### Implementation

LEAPH is implemented in python3.8.10 language (available at https://www.python.org/). The build-up process is done using jupyter-notebook v6.4.8 (68) while the final LEAPH-model and running software is a python3.8.10 script that can be used as stand-alone software or by executing a singularity v3.7 container (69) which includes all the required dependencies. EffectorComb was first implemented as R 4.3.2 (available at https://www.r-project.org/) script and then embedded into a shiny app (v1.8.0, available at https://shiny.posit.co/, R version) and provided within a singularity v3.7 container. Usage instructions and scripts are freely available at https://github.com/Plant-Net/LEAPH-EffectorComb.git.

## Results and Discussions

### A comprehensive collection of phytoplasmas effectors and features to describe their sequence characteristics

Careful selection of the training set and features to use for classification purposes are crucial in machine learning applications. By performing extensive literature and database mining, we collected 184 protein sequences from 53 phytoplasmas species that compose the “positive dataset”. The “negative dataset” is composed of 295 proteins whose function is not related to the known effectors’ activity and/or no interactions with host plant proteins are reported. (Figure 1a, Additional Table 1.1 and 1.2, refers to methods for further details). Intending to build a classifier for the prediction of novel effectors, we calculated 30 features to represent the salient characteristics of their sequences (Additional Table 3, please refer to methods for further details). We included eight features that describe the protein sequence properties and the mode of secretion, 12 features resuming the presence of protein domains important for plant invasion and infection and Ten features relating to the presence of characteristic sequence motifs found by MOnSTER (56). To inspect whether these features are discriminant of putative effector properties, we plotted the distribution of each, for the positive and negative datasets, respectively (Additional Figure 1). Twenty-four features show a significantly different distribution (Mann-Whitney test, p-value<0.05) by comparing the two datasets. However, we decided to include all features because non-significantly different features can be specific to a small class of putative effectors, too small to reach a statistical significance when including all the effector proteins together. Overall, both the datasets and the features are suitable for setting up a classifier based on machine learning models.

### The four classifiers included in LEAPH captured distinct features to predict pathogenicity proteins

We developed a novel ensemble learning classifier to predict pathogenicity protein candidates within phytoplasmas proteomes. Because of the wide variety of hosts and infection symptoms, we expect phytoplasmas’ effectors to have different characteristics, thus we reasoned that different learning models would be able to capture diverse properties yielding a more comprehensive prediction. Therefore, we used an ensemble learning composed of two tree-based algorithms, namely random forest and XGBoost (57,58), and two naive Bayes classifiers including a Gaussian and a Multinomial model (59,60). We fed the four classifiers with our training dataset and measured their performances on the test set (refer to methods for dataset construction and Additional Table 4 for models’ parameters). Due to the small size of the datasets, we performed five-fold cross-validation. Overall, the four models showed very good and comparable performance both within the cross-validation folds and between methods, ranging from 95% to 99% for the four measures used: precision, recall, accuracy and F1 score (Additional Table 6 and Additional Figure 2). This result supports our choice to include the four models to predict candidate effector proteins.

**Figure 2:**
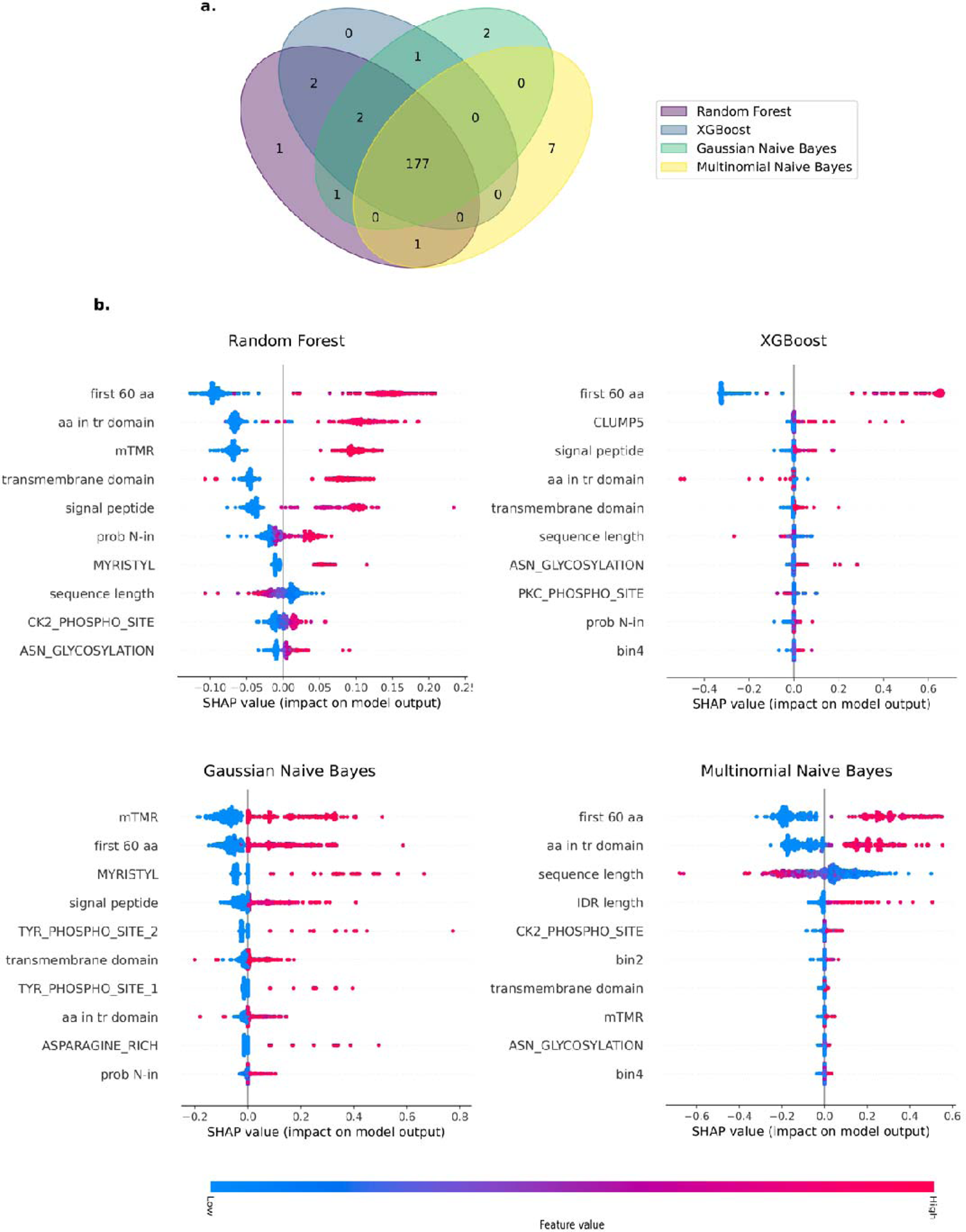
LEAPH and SHAP results on the training dataset. **(a)** Venn diagram representing the agreement of prediction on the training set from the four models included in LEAPH, namely: Random Forest model (purple ellipse), XGBoost (dark blue ellipse), Gaussian Naive Bayes model (aquamarine ellipse) and Multinomial Naive Bayes model (yellow ellipse). **(b)** the first ten features in terms of SHAP-value importance for each model included in LEAPH. Refer to Additional Figure 3 for the complete representation of SHAP importance.

The best performing model was selected for each classifier method and included in the ensemble approach that we call LEAPH (ensemb**L**e model for **E**ffector cl**A**ssification in **PH**ytoplasmas). To each candidate pathogenicity protein predicted by LEAPH, a score ranging from one to four is associated, corresponding to the number of classifiers that agree on the outcome.

In Figure 2a we report the Venn diagram showing the agreement of the prediction obtained by the four models. The majority of candidate effectors were predicted by the four models (96%), as expected. Interestingly, the two naive Bayes classifiers, and mainly the Multinomial model, were able to identify putative effectors that were not identified by the two tree-based models, supporting our hypothesis that different classifiers might capture different patterns of candidate pathogenicity proteins. To further investigate this direction, we calculated the feature importance for each classifier by using the SHAP algorithm (63), (SHapley Additive exPlanations) which is a game theory approach to explain the output of any machine learning model. Indeed, as shown in Figure 2b and Additional Figure 3, we observe a quite different spectrum of features with the highest contribution scores depending on the classifier. While the feature describing the first 60 amino acids of the protein sequence is found among the top two for all methods, the other features seem to contribute differently, depending on the classifier. In particular, we observe that only for the multinomial naive Bayes, the sequence length has a high impact on the model output. On the other hand, for the Gaussian naive Bayes model, we found the mispredicted transmembrane regions (mTMR) as the most contributing to this model. Altogether these results show that the four classifiers included in LEAPH can identify potential pathogenicity proteins with different characteristics and confer good performances on the test dataset.

**Figure 3:**
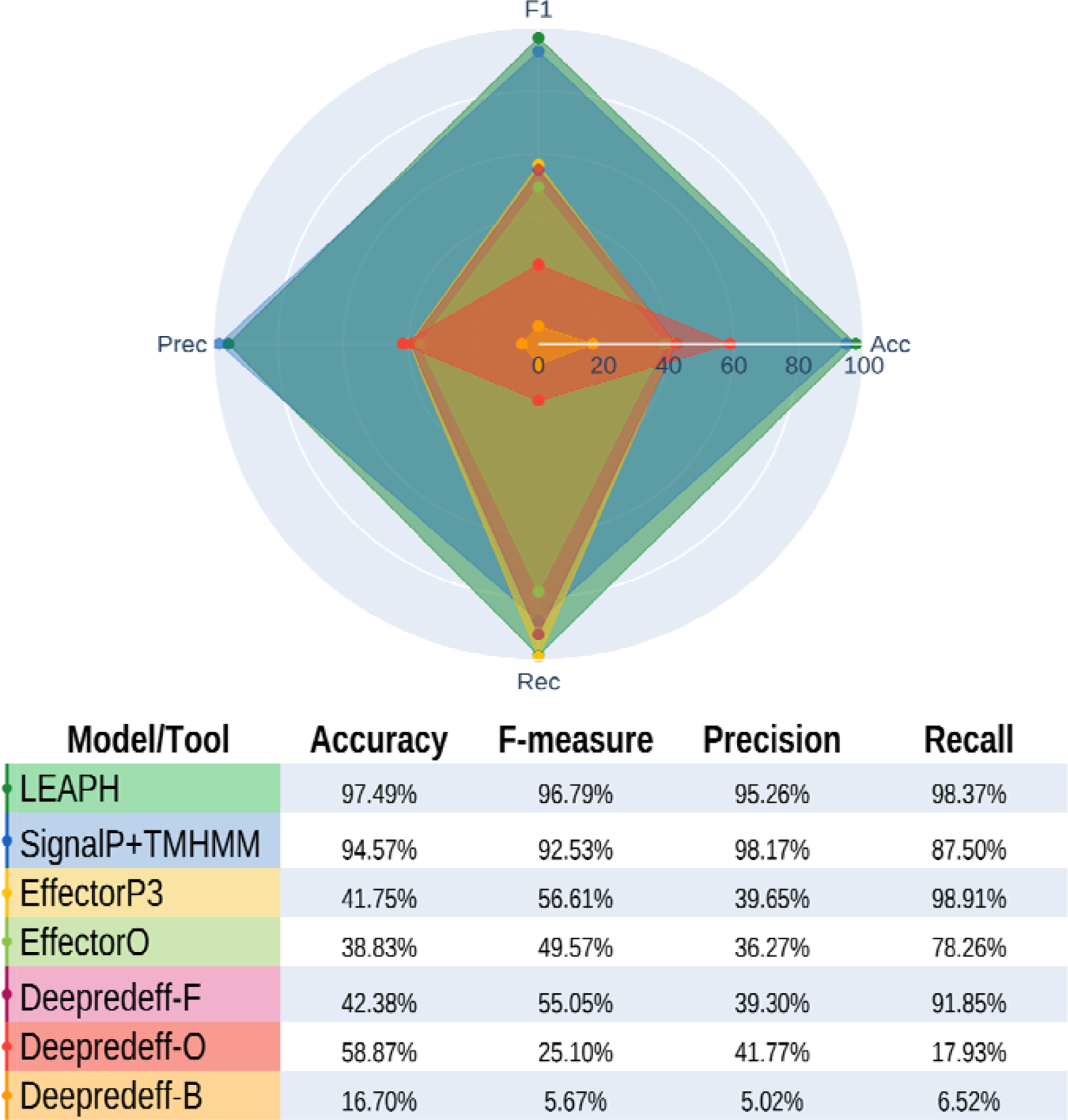
Performances of LEAPH and SOTA tools for effector prediction. Performances of each tested state-of-the-art tool compared to LEAPH performance. For each tool is reported the % of Accuracy, F-measure, Precision and Recall after the application on the training set.

### LEAPH outperformed other methods regarding the prediction of effectors in phytoplasmas

We compared the performances of LEAPH with four classifiers available for effector predictors, including SignalP4.1 (51) in combination, or not, with TMHMM2.0 (52) which is the only conventional method to predict putative effectors in phytoplasmas and EffectorO (14), EffectorP3.0 (17), the three models for Deepredeff (18) that are built to predict putative effectors in other species (Additional table 7 and Figure 3). LEAPH outperformed all the other methods by considering the four metrics used in this study, achieving a recall of 98.4%, thus predicting only 3 false negatives. In this context, the minimization of false negatives is crucial, to minimize the neglecting of putative true effectors. Importantly, LEAPH also reached good precision (95.3%), thus implying a low number of false positives as well (9 in total). Although experimental validation is capital to validate putative candidate effectors predicted by computational methods, the low number of both false negatives and false positives increases the confidence in the prediction and will permit a more successful experimental validation.

EffectorP3.0, EffectorO and Deepredeff-fungi showed quite good recall but very low values for the other metrics. This is due to the high number of false positives identified by those methods that lower the accuracy (ranging from 38.8 to 42.4%) and the precision (between 36.3 and 39.6 %). On the other hand, EffectorP3.0 and Deepredeff-fungi do not identify many false negatives with a recall of 98.9% and 91.8%, respectively. Lower performances are obtained for EffectorO with a recall of 78.3%. Deepredeff-oomycetes and Deepredeff-bacteria showed very poor performances. Altogether these results highlight the importance of developing a method tailored for phytoplasmas to identify *bona fide* candidate effectors and rule out EffectorP3.0, EffectorO and Deepredeff as tools to predict putative effectors in phytoplasmas.

The three combinations of SignalP4.1 with the TMHMM2.0, namely: SignalP4.1, SignalP4.1+TMHMM2.0, and SignalP4.1/TMHMM2.0 (in Additional Table 7), achieved comparable performances mainly in terms of accuracy concerning LEAPH. Despite that these tools are included among the features considered by LEAPH, their recalls are lower than the value achieved by our tool, meaning that these methods retrieve a higher number of false negatives, thus missing potential *bona fide* effector candidates. This can be explained by the fact that the combination of SignalP4.1 and TMHMM2.0 by definition can identify only putative classically secreted effectors, while there is a growing body of evidence supporting other ways of secretion of effectors not exhibiting the signal peptide, the so-called non-classically secreted effectors (41). We hypothesize that these false negatives can be potential non-classically secreted pathogenicity proteins that are captured by LEAPH only thanks to the ensemble of the four classifiers.

### LEAPH predicts classically, classically-like and non-classically secreted putative pathogenicity proteins from 13 phytoplasma proteomes

We selected 13 phytoplasma proteomes from 10 different species with different characteristics in terms of the number of proteins, 16S group, type of symptoms and type and number of hosts (Additional Table 8 and Figure 4). On average LEAPH predicts as putative pathogenicity proteins around 30% +/- 2.7 of the proteomes except for three strains: CaPhoenicium_SA213, CaPhoenicium_ChiP and CaPoryzae_NGS-S10 for which we have found 22.63%, 27.65% and 28.32%, respectively (figure 4a).

**Figure 4:**
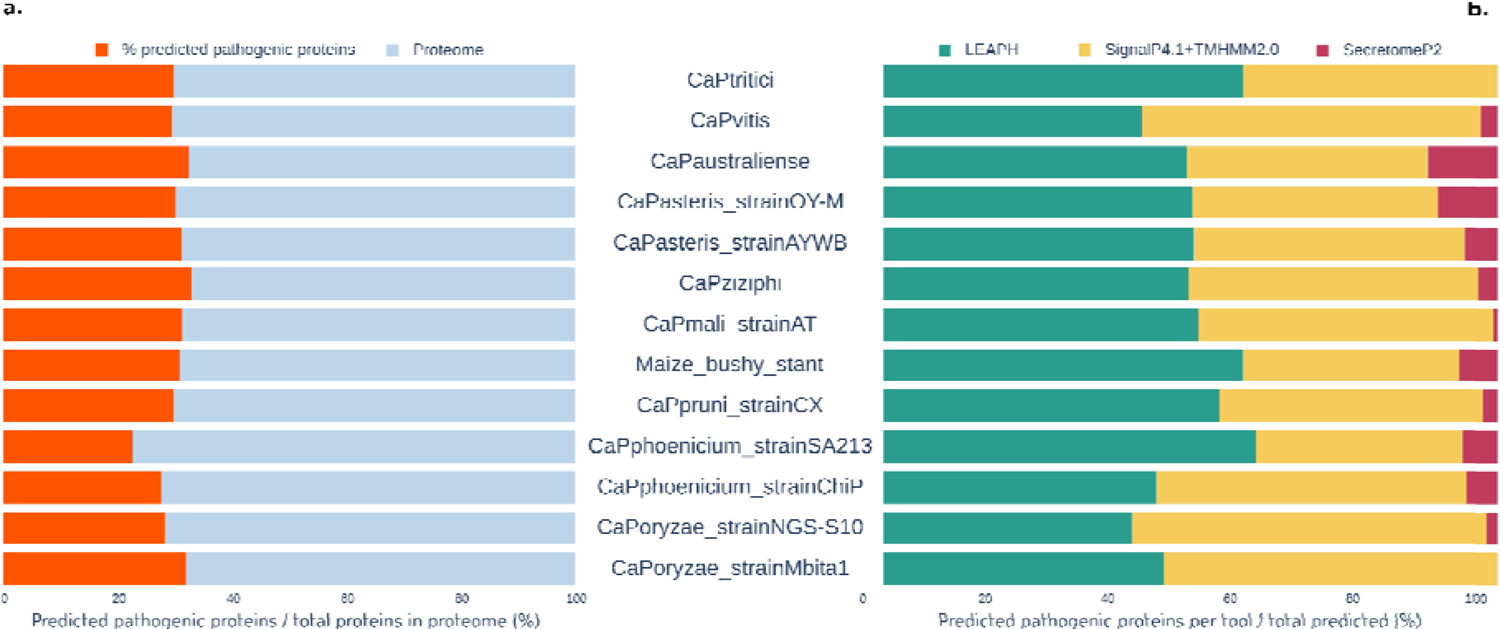
Putative pathogenicity proteins predicted by LEAPH from 13 phytoplasma proteomes. **(a)** Percentage of putative pathogenicity proteins predicted by LEAPH (orange bars) compared to the total percentage of proteins (grey bars) for the corresponding phytoplasma proteome. **(b) Comparison of t**he proportion of proteins predicted only by LEAH (green bars), predicted by the SignalP+TMHMM method and LEAPH (yellow bars) or by SectetomeP2.0 and LEAPH (magenta bars).

To characterize the properties of these putative pathogenicity proteins, we compared them with the prediction of SignalP4.1+TMHMM2.0 and SecretomeP2.0 (64). SignalP4.1+TMHMM2.0 is used to identify proteins having the signal peptide, usually present in classically secreted proteins, whereas SecretomeP2.0 predicts non-classically secreted proteins. While the first is currently used to predict putative effectors, this strategy can predict only classically secreted effectors. Importantly both methods predict whether the protein is secreted or not without information on its pathogenicity. Depending on the species, LEAPH predicts between 40 and 61% of putative effector proteins that were not identified by the other two methods (figure 4b). Between 33% and 58% of LEAPH putative effectors show also the signal peptide and the mispredicted transmembrane region identified by TMHMM2.0 (mTMR), thus implying that they are classically secreted. These results pinpoint that LEAPH, despite that it was trained on a dataset composed mainly of classically secreted effector proteins, can capture characteristics to identify potential effectors beyond the classically secreted ones. On the other hand, very few predicted effectors by LEAPH were also predicted by SecretomeP2.0. This was expected because the model used to run SecretomeP2.0 was suitable for gram-negative bacteria, but no model was available for gram-positive. According to Gao et al., (70) SecretomeP2.0 underestimates non-classically secreted proteins when using the gram-negative bacteria model for gram-positive ones. Interestingly, between 77% and 89% of putative effectors predicted exclusively by LEAPH contain only the mTMR (without the signal peptide), supporting the hypothesis that phytoplasmas effectors might have a peculiar signal peptide located into the so-called Sequence-Variable Mosaics regions of some proteins (71).

Finally, we checked whether the known validated phytoplasmas effectors were correctly classified by LEAPH. Eleven classically secreted effectors were identified and validated by previous studies: two SAP11-like proteins in two different species (‘*Ca*. P. asteris’ strain AYWB and ‘*Ca*. P. mali’ strain AT (36,46)), SAP54 (‘*Ca*. P asteris’ strain AYWB (36)), SAP05 (‘*Ca*. P. asteris’ strain AYWB (36)), PHYL1 (‘*Ca*. P. asteris’ strain OY-M (40)), TENGU (‘*Ca*. P. asteris’ strain OY-M (35)), SWP1, SWP11, SWP12, SWP16 and SWP21 (‘*Ca*. P. tritici’ (72–75)); and six non-classically secreted effectors in the strain ‘*Ca*. P. Ziziphi’: ncSecP3, ncSecP12, ncSecP14, ncSecP22, ncSecP9 and ncSecP16 (70) (Additional Table 9). Remarkably, not only LEAPH could correctly identify the eleven classically secreted, but also four out of six non-classically secreted validated effectors (ncSecP3, ncSecP12, ncSecP14, ncSecP22). This result strengthens the potentiality of LEAPH to identify putative pathogenicity proteins independently of their type of secretion.

### Description of the effector proteins landscape predicted by LEAPH

To further characterize the putative pathogenicity proteins predicted by LEAPH, we performed a principal component analysis (PCA) on protein features (see methods). In Figure 5 we observe three distinct groups. To understand whether these groups could represent some property of protein sequences, we colored according to the type of secretion. We defined as “classically” secreted, those proteins in which both signal peptide and mTMR are predicted; “classically-like” secreted, those having only the prediction for the mTMR; “non-classically” secreted, those proteins in which neither signal peptide nor mTMRs are predicted (Figure 5a). The three groups identified by the PCA respectively follow the three secretion modalities. Thus, we checked where the validated effectors are located in the putative pathogenicity proteins landscape identified by the PCA (Figure 5b). The eleven classically secreted validated effectors (two SAP11 in two different strains, SAP54, SAP05, PHYL1, TENGU, SWP1, SWP11, SWP12, SWP16 and SWP21) were localized in the cluster of classically secreted as expected. Surprisingly, among the four non-classically secreted validated effectors, only two were found in the correspondent cluster, namely ncSecP3 and ncSecP12. Unexpectedly, by inspecting the protein sequences, we found that ncSecP14 contains the signal peptide and the mTMR and ncSecP22 only the mTMR, thus confirming their localisation on the PCA. Although further investigations will be needed to assess the final class of these effectors, this is out of the scope of the present work. Altogether these findings allowed us to define three groups of putative pathogenicity proteins predicted by LEAPH as the classically, the classically-like, and the non-classically secreted.

**Figure 5:**
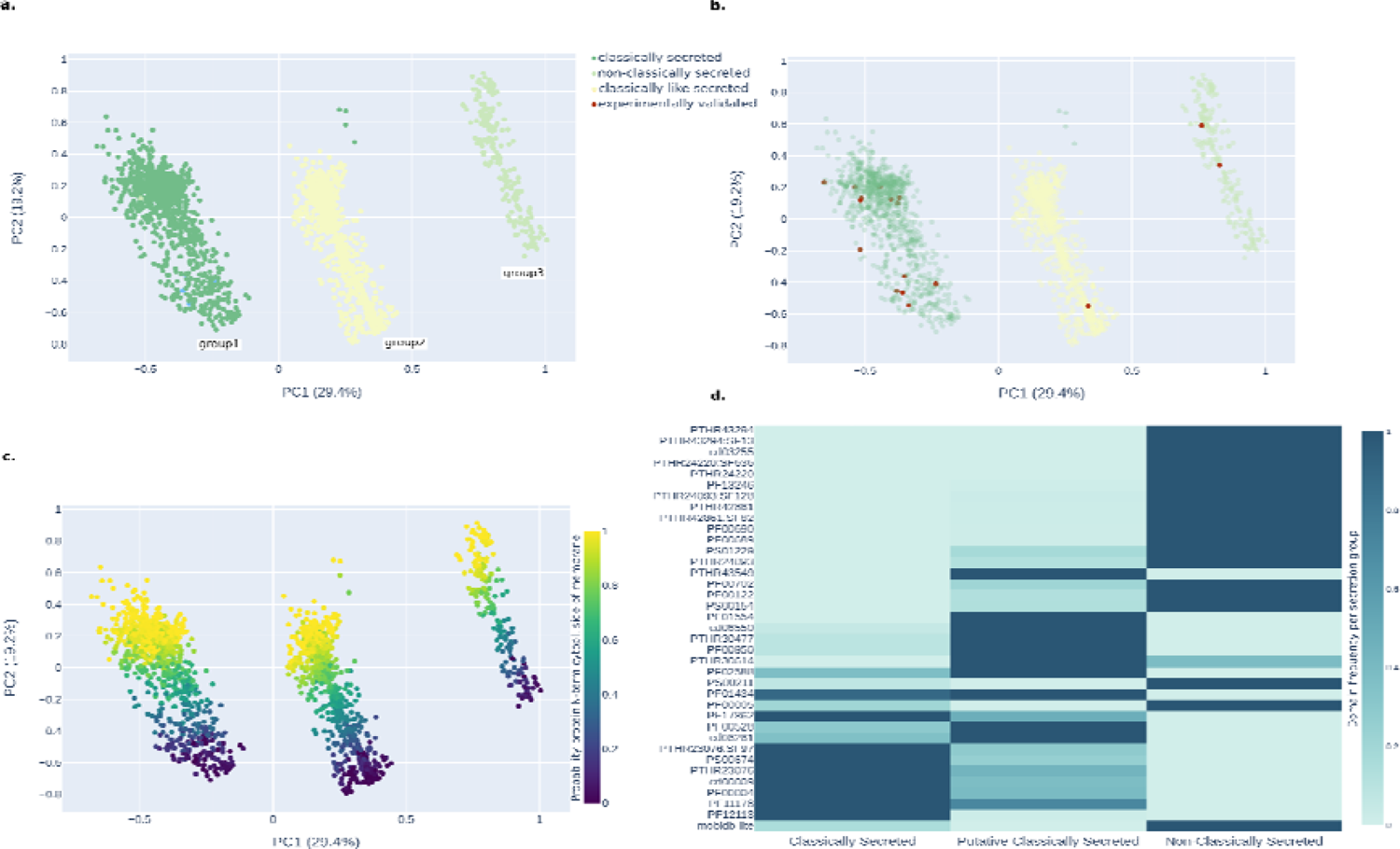
Description of the effector proteins landscape predicted by LEAPH. **(a)** PCA of predicted pathogenicity proteins by LEAPH from 13 phytoplasma proteomes coloured by secretion methodology; group1 (dark green points) predicted by SignalP4.1 to bear a signal peptide (classically secreted), group2 (light yellow) contain a mispredicted transmembrane region in the first 60 amino acids of the sequence (mTMR) from TMHMM2.0 and no prediction of signal peptide by SignalP4.1 (classically-like secreted) and group3 (light green points) do not have the signal peptide and the mTMR (non-classically secreted). **(b)** Same as (a) with the position of the 15 experimentally validated effectors from the literature (dark red dots). **(c)** same as (a) coloured by the probability that the N-terminal region of the protein is in the inner part of the membrane. **(d)** Depict the abundance of each protein domain predicted by InterProScan in each PCA group.

Then, we mapped on the three PCA groups other properties. Interestingly, inspecting the second principal component, we noticed that the putative pathogenicity proteins found by LEAPH are stratified by the probability for the N-terminal to have a cytoplasm location, as shown in Figure 5c. When colouring by species, we observed that there are no specific clusters, as expected, meaning that pathogenicity protein properties are not species-specific (Additional Figure 4a). In Additional Figure 4b we can observe that proteins are stratified by sequence length. Surprisingly we observe that few proteins from the classically secreted and non-classically secreted groups have a size larger than expected. Thus, we mapped these proteins on the sequence length distribution of the respective proteomes to see whether these sizes could be potential outliers or not (Additional Figure 4c). We observed that very few putative pathogenicity proteins, mainly predicted as classically secreted, map on the tails of these distributions.

To investigate other possible peculiar properties of LEAPH-predicted putative pathogenicity proteins, we used InterProScan5 (66) to predict protein domains in the three classes of secretion. We considered only domains with an occurrence of at least 1% for at least one of the secretion classes, for a total of 37 protein domains. As shown in Figure 5d, classically secreted putative pathogenicity proteins mainly possess domains characteristic of both SVM-proteins, AAA+ ATPases and FtsH proteins. Others distinguish SVM or Sequence Variable Mobile proteins for the presence of a modified secretion signal (71)). Recently, it has been demonstrated that the AAA+ and Ftsh domains have a role in pathogenicity for ‘*Ca.* P. mali’ (76,77). Classically-like secreted candidate pathogenicity proteins, on the other hand, are mostly characterized by periplasmic binding proteins or ABC-transporters and only slightly enriched in AAA+ and Ftsh domains. Although ABC-transporters are found ubiquitously in eukaryotes and prokaryotes, they play a crucial role in pathogenesis and virulence in pathogen bacteria (78). There are different types of ABC transporters (ten according to y. Zeng and A.O. Charkowski (79)) and this might be linked to the presence of different ABC-related domains characterizing the non-classically secreted group of predicted pathogenicity proteins. In this class, ABC transporters are less frequent concerning the P-ATPases and especially to the intrinsically disordered regions. P-ATPase, particularly copper exporters P-type ATPases are reported to play a major role in the virulence of diverse pathogenic bacteria even if the underlying mechanisms remain partially understood (80–84)). These findings suggest that putative pathogenicity proteins are distinguished by the secretion mode and other characteristics related to virulence properties embedded in the sequence composition.

### SOM clustering allowed us to build a pathogenicity proteins reference map for phytoplasmas

To study whether other properties are characteristics of sub-classes of pathogenicity proteins beyond the type of secretion, we performed a clustering analysis on the LEAPH-predicted proteins using the SOM model (67). This analysis allowed us to create a 2D map composed of a 10×10 lattice where each hexagonal cell is characterized by a peculiar combination of features and closer cells have more similar properties than farther cells. Thus, the 2093 putative pathogenicity proteins, including the 15 validated effectors, identified by LEAPH across 13 phytoplasma proteomes are distributed into the map and associated to a specific hexagonal cell according to their sequence properties. Consistently with PCA analysis, the two main properties that stratify the hexagons in the lattice are the signal peptide and the mTMR on the x-axis, and the probability for the N-terminal to have a cytoplasmic location on the y-axis (Figure 6a). Therefore, we can visualize on the map the hexagons that correspond to the three groups found by the PCA analysis: 49 hexagons for classically secreted, 31 for classically-like and seven for non-classically secreted putative pathogenicity predicted proteins. Furthermore, eight hexagons that contain proteins from both groups of classically-like and non-classically secreted and five empty hexagons can be identified on the map. We foresee that with the advancement of phytoplasma research, the availability of proteomes will increase as long as the identification of new putative pathogenicity proteins and these empty hexagons will be possibly filled by proteins showing new properties combinations.

**Figure 6:**
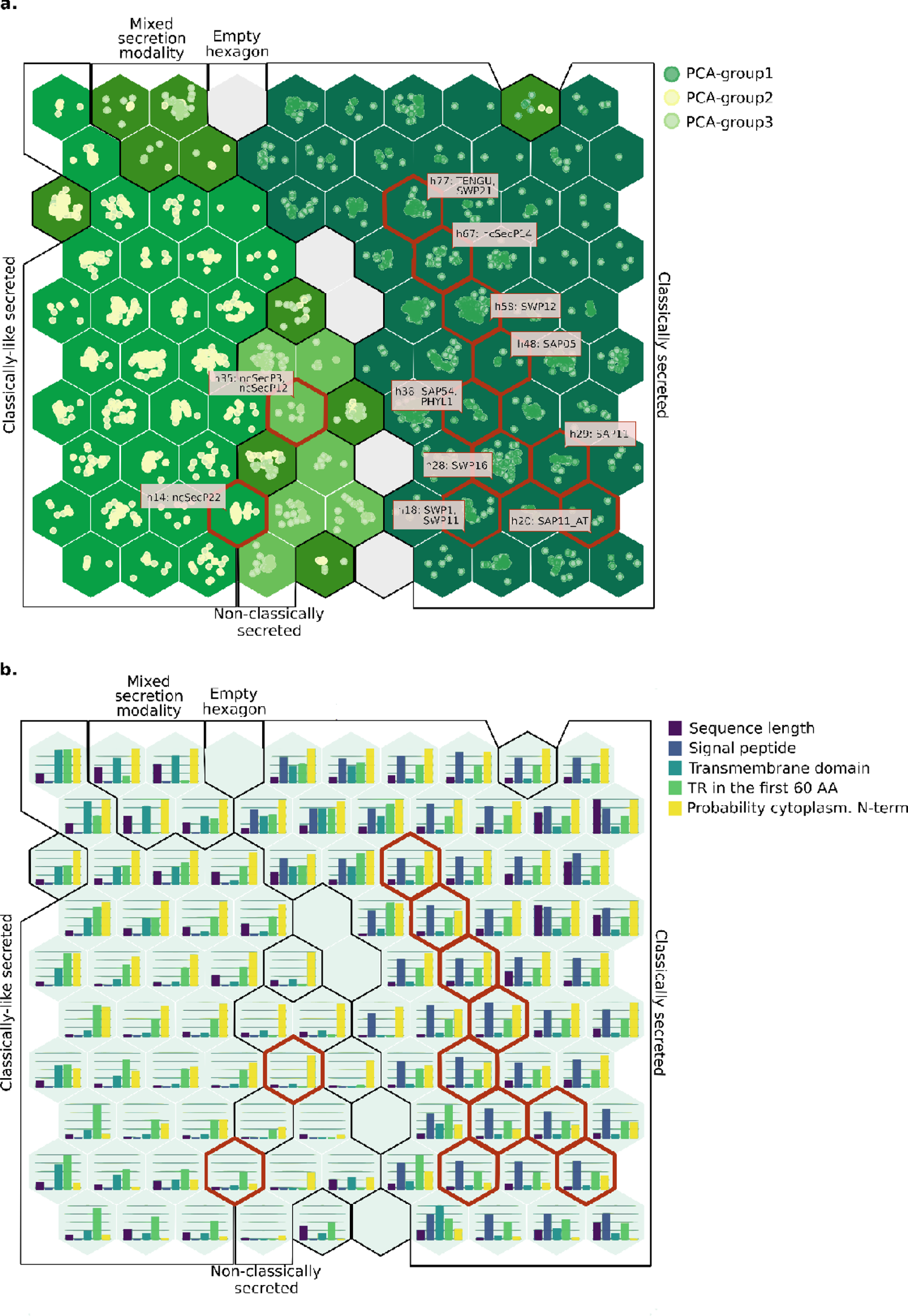
Pathogenicity proteins reference map for phytoplasmas. **(a)** LEAPH predicted proteins projected on a Self-Organizing Map (SOM) and coloured by their group of secretion: classically secreted (dark green points and dark green hexagons), classically-like secreted (light yellow points and green hexagons), non-classically secreted (light green points and light green hexagons), mixed secretion modality (olive green hexagons). Dark red contours indicate the hexagons containing the validated effectors from the literature. Empty grey hexagons represent profiles of features that are not represented by the predicted proteins. **(b)** same as (a) showing the features abundance for each hexagon.

Interestingly, the 15 validated effectors are projected in nine different and close hexagons (including ncSecP14 proposed by Gao et al., (70) as a non-classically secreted but having the signal peptide as discussed in previous paragraphs) (Figure 6a). As expected the two homologous phyllogen proteins (PHYL1 and SAP54) are in the same hexagon (h38). Similarly, SWP21 and TENGU are in the same hexagon (h77). The SAP11-like proteins do not fall in the same hexagon but in hexagons next to each other, suggesting more variability in the sequence properties of proteins in this class depending on the phytoplasma species. Intriguingly, SWP1 and SWP11, which have different roles in the disease development (72), are in the same hexagon (h18). Interestingly, both interact with the TEOSINTE BRANCHED 1/ CYCLOIDEA/ PROLIFERATING CELL FACTOR 1 and 2 (TCP) transcription factors, thus suggesting shared sequence properties to enhance the interaction with similar partners to further investigate (85). The two non-classically secreted validated effectors (ncSecP12 and ncSecP3) are mapped in the same hexagon (h35) also suggesting very similar sequence properties. Importantly, LEAPH predicted new putative pathogenicity proteins that are mapped in the same hexagons as the biologically validated effectors offering novel potential effector candidates with similar sequence characteristics to be experimentally validated.

### EffectorComb: a shiny app to inspect the pathogenicity proteins map

To further inspect other properties and characteristics of the sequences contained in the pathogenicity protein map, we have developed a shiny application, called EffectorComb, enabling us to show which features are enriched in each hexagon (Additional Figure 5). For instance, we notice that the enrichment of phosphorylation sites is mainly present in hexagons collecting classically secreted putative pathogenicity proteins. Specifically, hexagon h70 is characterized by a high presence of the domain cAMP- and cGMP-dependent protein kinase phosphorylation site. Similarly, the Protein kinase C phosphorylation site and Casein kinase II phosphorylation site were found predominantly in the h90. Tyrosine kinase phosphorylation sites 1 and 2 were also found enriched in h25, h99, h100 and h48. Protein phosphorylation is a widely used mechanism in bacteria to adapt to changes in their environment (where the conditions can alter rapidly), but it is also used for intercellular communication. The N-glycosylation site, specific to the consensus sequence Asn-Xaa-Ser/Thr, was found enriched in both classically-like (h74) and classically secreted (h89) candidate pathogenicity proteins. Bacteria have evolved diverse glycosylation systems for pathogenesis. Bacterial glycosylation not only allows adhesion to the host cell but also functions to modulate crucial host cellular processes (86). Notably, glycosylation has been reported as an important process in the interaction between the ‘Flavescence dorée’ phytoplasma and its insect vector (87), but also of phytoplasmas strains and host plants (88,89). Interestingly the hexagon h74 is also characterized by the presence of the N-myristoylation site. N-myristoylation is an irreversible protein lipidation, in plants, it is recognized as a major modification and concerns nearly 2% of all plant proteins (90,91). Precisely, attachment of a myristoyl group increases specific protein–protein interactions leading to the subcellular localization of myristoylated proteins with their signaling partners (92). Intriguingly, Medina-Puche et al. (93) identified novel mechanisms involving protein relocation due to the presence of the N-myristoylation site from the plasma membrane to chloroplasts that are utilized in the arms race between plants and pathogens to regulate the other side in favor of defense responses in plants and the pathogenesis of pests. Furthermore, amidation sites were found mainly in the h10 and h40. C-terminal amidation reduces the overall charge of a peptide; therefore, its overall solubility might decrease. Modifications at the C-terminal have already been shown to have consequences on the pathogenicity of phytoplasmas, for instance, deletions at the C-terminal of the effector SWP1 impaired the induction of witches’ broom (94). The enrichment of this domain might suggest that changes in protein solubility are important for phytoplasmas’ effectors and might reflect distinct functions depending also on the different hosts (insect/plant). The leucine zipper domain is found only in the classically secreted putative pathogenicity proteins lying in h19. This pattern is present in many gene regulatory proteins making this cluster very interesting for further investigation (95). Finally, the ‘RGD’ tripeptide that has been shown to play a role in cell adhesion (96), is abundant in both classically (h82,h98) and non-classically (h1, h17, h25) predicted pathogenicity proteins. Overall, the SOM map allowed us to identify distinct classes of putative pathogenicity proteins with peculiar properties probably linked to their function and type of interaction with the plant and/or insect.

## Conclusions

The few known biologically validated effectors in phytoplasma are implicated in a huge spectrum of different functions, going from the regulation of plant morphogenesis to attracting insect vectors. They possess a variety of strategies to manipulate the host plants. Therefore, it is unlikely for phytoplasma effectors to be described by a few common characteristics. However, the most widely used method to select pathogenicity candidates for phytoplasma is to screen protein sequences and look only for the presence of the signal peptide. Leveraging four machine learning classifiers coupled with a novel voting score for assessing the prediction score, we have developed LEAPH specifically conceived for phytoplasmas’ effector candidate prediction. We have shown that LEAPH can predict putative effector candidates with high reliability and outperform existing tools not adapted for phytoplasma. A major advantage of LEAPH consists in its ability to predict candidate pathogenicity proteins independently by the presence of the signal peptide and with many different sequence characteristics. Furthermore, as our knowledge of effector proteins increases, LEAPH can be refined by constituting a novel training dataset to re-train the models to improve their performances.

The application of LEAPH on 13 phytoplasma proteomes led to the identification of 2093 putative candidates showing sequence characteristics linked to virulence and pathogenicity. To investigate the properties of those proteins we used the SOM algorithm and obtained the first pathogenicity proteins map for this pathogen. To allow easy exploration of the pathogenicity protein map, we have developed an easy-to-use shiny application called EffectorComb. To the best of our knowledge, this is the first time that a comprehensive characterisation of putative pathogenicity proteins has been provided for phytoplasma. The use of this map goes beyond the identification of groups of putative pathogenicity proteins having similar properties but can be used as a reference map to project new predicted pathogenicity proteins from other proteomes in a predictive perspective. Therefore, when using this map as a reference to analyze a new sample, the positioning of each predicted pathogenicity protein of a new proteome on the 2D plane can be used to assess its pathogenicity properties and overall sequence characteristics. The pathogenicity proteins map can be used also to accelerate the choice of novel candidates for biological validation for instance by focusing on proteins in the same group of known validated effectors or on groups showing peculiar sequence properties of interest.

Overall, LEAPH and EffectorComb are valuable tools to improve our understanding of effectors in plant–phytoplasma interactions. Finding novel experimentally validated effectors can be used to set up novel methods to improve plant resistance to this devastating pest.

## Availability and requirements

The datasets supporting the conclusions of this article are included within the article and its additional files.

- Project name: LEAPH-EffectorComb
- Project home page: https://github.com/Plant-Net/LEAPH-EffectorComb.git.
- Archived version: 10.5281/zenodo.10276703
- Operating system(s): Platform independent but software requirements to be fulfilled
- Programming language: Python3.8, R for shiny-app
- Other requirements: biopython, pandas, joblib, SignalP4.1,TMHMM2.0, MobiDB-lite3.0, Prosite1.86, singularity3.7 (at least)
- License: e.g. GNU GPL

## Declarations

### Availability of data and materials

LEAPH, along with training sequences, feature tables, and Shiny App are available on GitHub at: https://github.com/Plant-Net/LEAPH-EffectorComb.git. The pre-trained LEAPH and the Shiny App are also available as Singularity 3.7 containers.

### Authors’ contributions

GC developed LEAPH and EffectorComb tools, prepared the datasets, performed all analyses and interpreted the results, drafted the manuscript, AC participated in study and tool conception, data interpretation and manuscript revision, HS participated in data interpretation and manuscript revision, KJ participated in data interpretation and manuscript revision, CD participated in data interpretation and manuscript revision, MM participated in data interpretation and study conception, revised the manuscript, SB conceived the study and tools, interpreted the results, drafted the manuscript. All authors revised and approved the manuscript.

## Acknowledgements

This work was supported by ELIXIR-IT (ELIXIR-IIB, ELIXIR-ITA, Elixir Italy, ELIXIR IIB, ELIXIR IT), the Italian research infrastructure for life-science data.

