## additional figures for "Definition of the effector landscape across 13 Phytoplasma proteomes with LEAPH and EffectorComb"

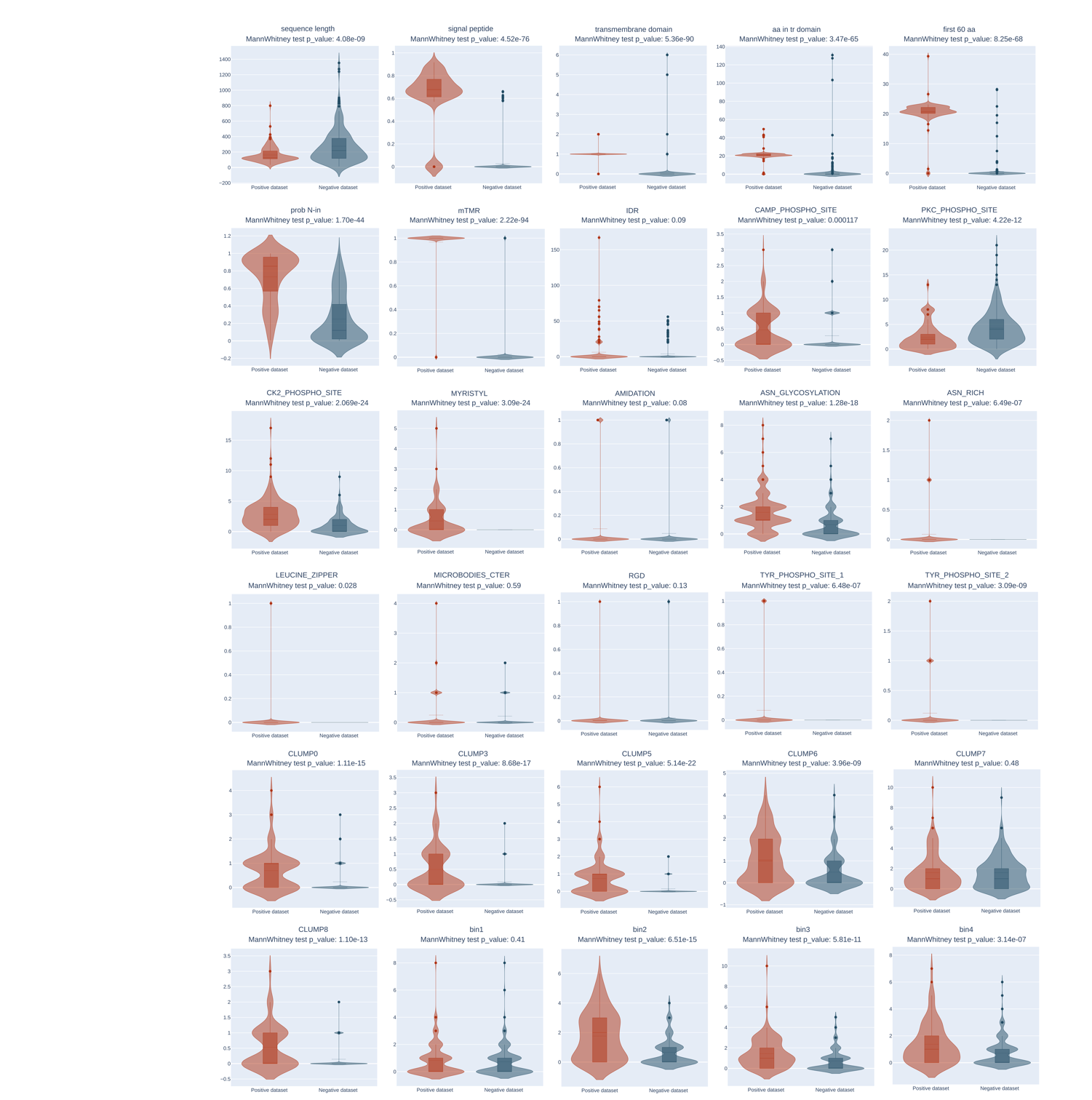

**Additional Figure 1: Features distribution for both positive and negative dataset.**

Distribution and corresponding p-value from Mann–Whitney Test on positive and negative dataset, for each of the 30 features included in the LEAPH model.

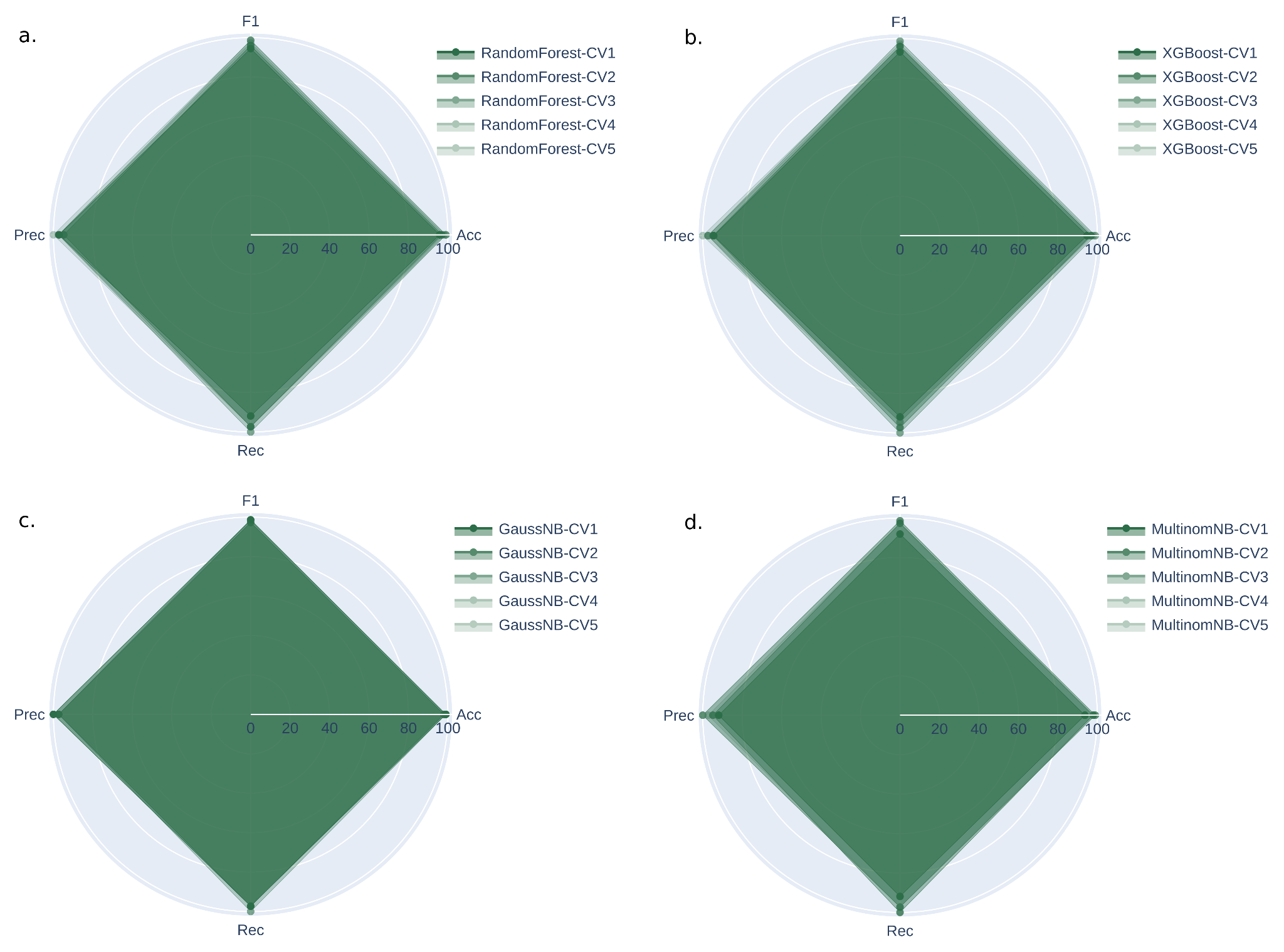

**Additional Figure 2: Cross-validation performances for each model included in LEAPH.**

**(a)** Shows performances of Random Forest model for each fold in the cross-validation process (5 fold in total). Performances are evaluated in terms of Accuracy, F-measure, Precision and Recall. Likewise **(b)** Shows performances in cross-validation for XGBoost, **(c)** for Gaussian Naive Bayes and **(d)** for Multinomial Naive Bayes.

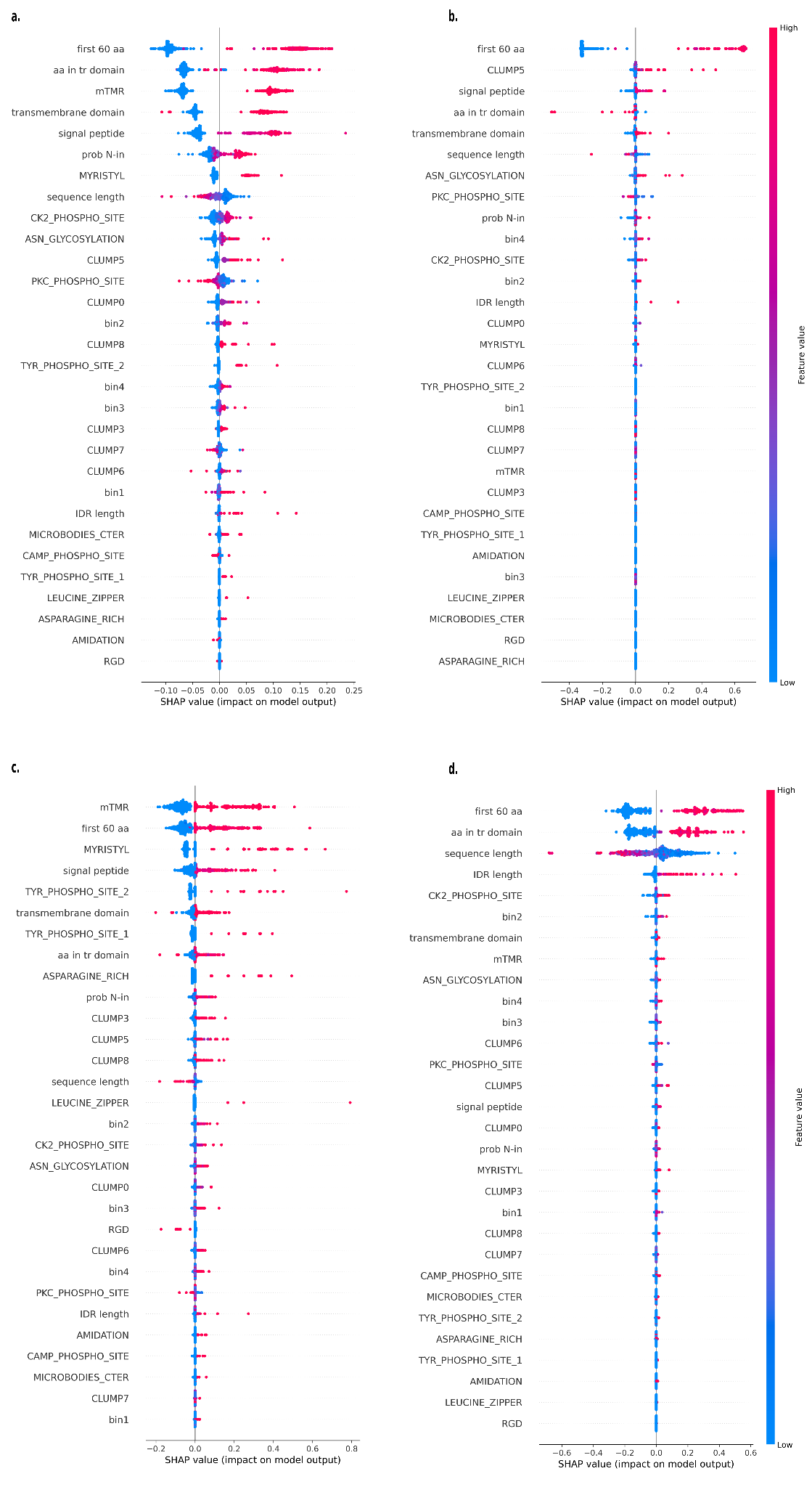

**Additional Figure 3: SHAP importance values for each feature for each model in LEAPH.**

**(a)** Represents the SHAP importance values for each of the 30 features included in the LEAPH model, for the Random Forest best model. Likewise **(b)** represents the importance of features in XGBoost, **(c)** the importance of features in Gaussian Naive Bayes and **(d)** show the importance of features in Multinomial Naive Bayes model.

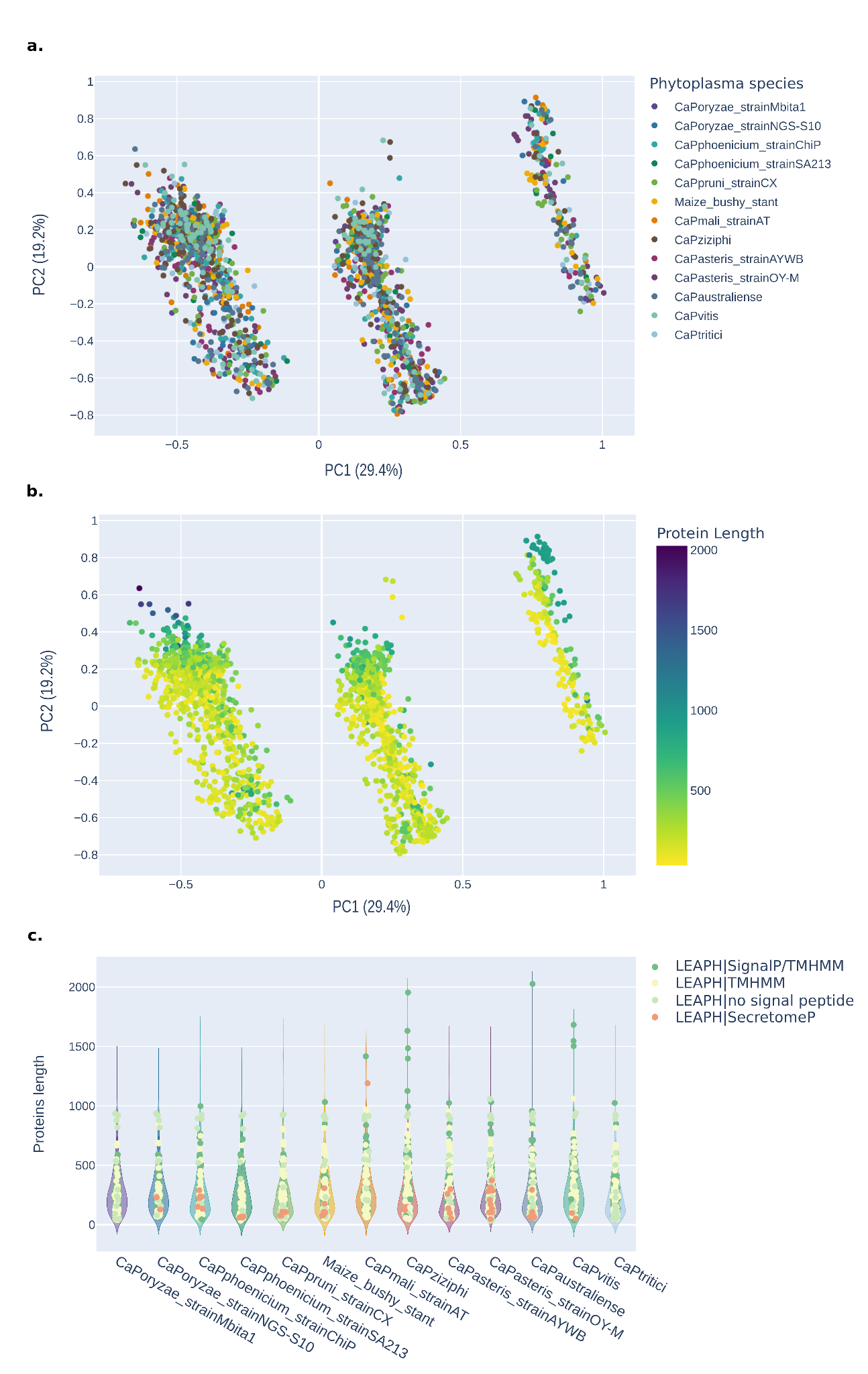

**Additional Figure 4: PCA of LEAPH predicted pathogenicity proteins and protein length distribution per proteome.**

**(a)** Shows the PCA of the predicted proteins by LEAPH, colored by the belonging species. **(b)** Shows the PCA of LEAPH predicted proteins colored by protein length. **(c)** Represents the protein length distribution of each of the 13 phytoplasma proteomes and the length of corresponding LEAPH predicted putative pathogenicity proteins. Dark green dots represent predicted proteins having a prediction of signal peptide from SignalP4.1 and a mispredicted transmembrane region from TMHMM2.0, light yellow dots represent predicted proteins having a mispredicted transmembrane region by TMHMM2.0, light green dots represent proteins having no prediction from the previous tools, dark pink dots represent proteins predicted by both LEAPH and SecretomeP2.

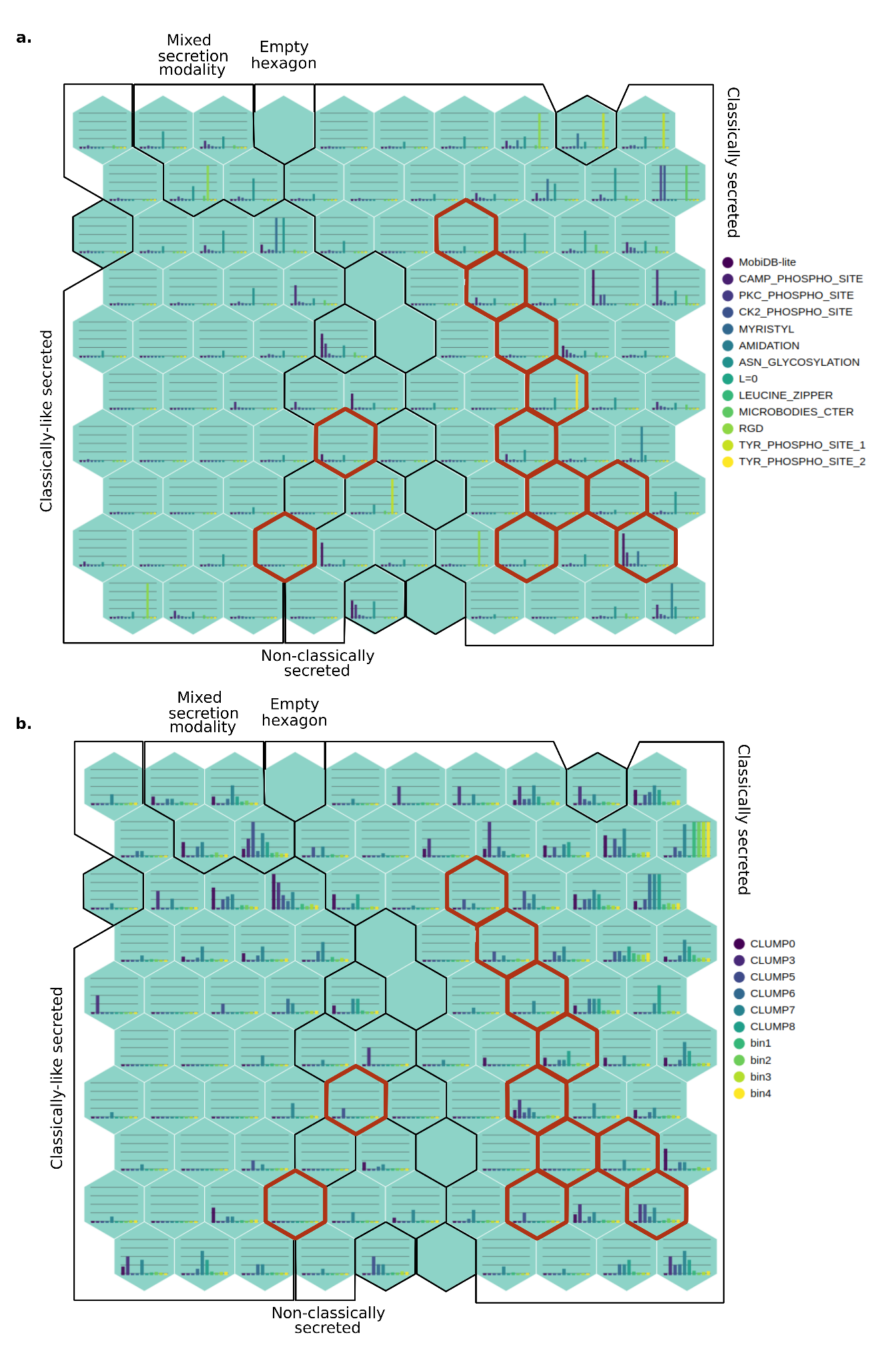

**Additional Figure 5: Self-Organizing Maps showing different set of features.**

**(a)** Shows the Self-Organizing Map (SOM) of LEAPH predicted putative pathogenicity proteins in terms of functional motif enrichment for each hexagon in the SOM. Similarly **(b)** represents the SOM of predicted proteins in terms of CLUMPs (from MOnSTER software) enrichment.
